# High within-tree leaf trait variation and its response to species diversity and soil nutrients

**DOI:** 10.1101/2023.03.08.531739

**Authors:** Andréa Davrinche, Anna Bittner, Helge Bruelheide, Georg Albert, W. Stanley Harpole, Sylvia Haider

## Abstract

- Leaf functional traits provide important insights into plants’ responses to different environments. Leaf traits have been increasingly studied within-species in the last decade, following the growing realisation that neglecting the intra-specific scale can result in misreading plants’ response to environmental change. However, while likely to lead to similar pitfalls, within-individual leaf traits are under-researched despite being the scale at which elementary interactions shape ecosystem processes.
- To address this critical lack of understanding at the local scale, we assessed leaf trait variation in a large biodiversity-ecosystem functioning experiment in subtropical China. We used optical spectroscopy to determine nine morphological and biochemical traits of >5800 leaves from 414 trees representing 14 species. We evaluated the relative importance of the intra-individual level for total leaf trait variation, and the interacting effect of two trait variation’s drivers, soil nutrient availability, and a local species richness gradient.
- Comparing the amount of trait variation at the between-species, between-individuals and intra-individual levels, we found that intra-individual variation accounted on average for >25% of total trait variation. Additionally, intra-individual variation was the most prominent component of intra-specific variation. We found partial support for positive effects of soil nutrient availability and species diversity on intra-individual trait variation, and a strong interdependence of both effects. Contrary to the amplifying effects we expected, trait variation increased with soil nutrient availability at intermediate diversity, but decreased at low and high diversity.
- Our findings quantify the relevance of intra-individual level for leaf trait variation, and expose a complex interaction between its drivers. In particular, interactive effects of soil nutrient availability and local species diversity on trait variation suggest responses ranging from alleviating competition to enhancing complementarity. Taken together, our work highlights the importance of integrating an intra-individual perspective to understand trait-based mechanisms in biodiversity-ecosystem functioning relationships.

## Introduction

Plant functional traits in general (Cadotte 2017; Liu et al. 2016; Reich, Walters, and Ellsworth 1997) and leaf functional traits specifically (I. J. Wright et al. 2004) can provide essential insights into ecosystem processes. Traits can, for example, inform how species interact with each other (Bastias et al. 2020; Kraft, Godoy, and Levine 2015), and how species contribute to ecosystem functioning (Ratcliffe et al. 2017; Mokany, Ash, and Roxburgh 2008b). In particular, analysing how traits vary with biotic and abiotic environmental conditions can help to understand those factors’ effects on the ecosystem (Trugman et al. 2019; Joswig et al. 2022).

It is generally assumed that plant traits vary the most between species, as opposed to within species (Garnier et al. 2001; McGill et al. 2006; Shipley et al. 2016). Hence, most studies have focused on between-species trait variation and described species by their mean trait values (Lavorel et al. 2008; Weiher et al. 2011), implying that trait variation within the same species (intra-specific trait variation) is negligible. However, such assumptions were critically challenged in the last decade (e.g. Albert et al., 2010; Fajardo & Piper, 2011; Hulshof & Swenson, 2010; Messier, McGill, & Lechowicz, 2010), and multiple studies argued to broaden this view (Westerband, Funk, and Barton 2021; Des Roches et al. 2018; Violle et al. 2012). In particular, recent work has raised awareness of the limitations resulting from ignoring intra-specific trait variation, such as underestimating functional diversity (Albert et al. 2012) or overlooking mechanisms affecting species coexistence (Hart, Schreiber, and Levine 2016). For example, Lepš *et al*. (2011) showed that intra-specific variability explained most of the community-level variation for some traits in response to different combinations of environmental conditions, and thus be dominant compared to, for example, species turnover alone. Ignoring intra-specific trait variation might therefore lead to an underestimation of the effects of environmental pressures on plant communities. As a result, interest in intra-specific trait variation gained considerable momentum, revealing that, on average, about a quarter of total variation can be explained by differences within species (Siefert et al. 2015), this proportion even reaching up to 40 % (Kattge et al. 2011).

In addition to substantial variation within species, traits can also vary greatly within an individual (i.e., intra-individual variation). So far, in plant functional ecology, only few studies have zoomed in from intra-specific trait variation to intra-individual trait variation (Umaña et al. 2018; Messier, McGill, and Lechowicz 2010; Auger and Shipley 2013; Kang et al. 2014), and several have suggested that intra-individual trait variation can even be greater than between-individuals trait variation (Herrera, Medrano, and Bazaga 2015; Kafuti et al. 2020). Ignoring the intra-individual scale of variation means neglecting an essential aspect of the answer to understanding the structure and functioning of plant communities. As similarly described by Messier et al. (2010) for intra-specific variation, trait variation of the individual is subject to assumptions on its importance that are the basis of approaches aiming at clarifying how and why species coexist and interact with each other and their environment. Hence, it is essential to identify the scales at which a large share of the trait variation occurs in order to apply ecological theories realistically, but also to study ecological processes at the very scale where they have a defining importance for the ecosystem – that is, where individuals interact. For example, Valdés-Correcher *et al*. (2021) showed that in natural forest stands, accounting for the intra-individual scale when explaining variation of leaf defence and herbivory-related traits enables a better assessment of trees’ genetic relatedness as an explanatory factor. As a result, ignoring intra-individual variation would occult the importance of genetic signals when explaining trees’ response to herbivory. However, in practice, studying leaf traits at the intra-individual level involves a high workload and associated costs, in addition to limitations due to small amounts of leaf biomass available for analysis. Because of these limitations, spectroscopy, a fast and non-destructive high-throughput method (Serbin et al. 2014; Burnett et al. 2021), has emerged as an advantageous tool for determining leaf traits, allowing the inclusion of thousands of samples to quantify intra-individual leaf trait variation (Proß et al. 2021; Davrinche and Haider 2021). Because of the low number of studies addressing intra-individual leaf trait variation, little is known about its drivers. However, trait variation at the intra-specific scale was found to be influenced by the abiotic environment (Lemke et al. 2015; Souza et al. 2018; Proß et al. 2021). For example, several experimental studies showed that high soil nutrient availability enhanced intra-specific trait variation (Lemke, Kolb, and Diekmann 2012; Helsen et al. 2017). In contrast, the relationship between resource availability and intra-individual trait variation has hardly been studied so far. Therefore, the question arises whether the results found at the intra-specific level are transferable to the intra-individual level.

In addition to abiotic drivers, biotic factors likely influence intra-individual traits and their variation. In particular, competition, a dominant driver of plant interactions (Connell 1983; A. Wright, Schnitzer, and Reich 2014), constitutes a strong interaction between individuals of the same species that require the same resources, and have similar uptake pathways (Grime 1973; Tilman 1982; Barabás, Michalska-Smith, and Allesina 2016). This can result in less resources available for each individual aboveground (e.g., light; Pretzsch, 2014) or belowground (e.g., nutrients; Pornon, Escaravage, & Lamaze, 2007). Inversely, diversity of co-existing species can decrease competition and promote the availability of resources through resource-use complementarity (Loreau and Hector 2001; Cardinale et al. 2007; Barabás, Michalska-Smith, and Allesina 2016). For example, several studies showed higher soil total phosphorus supply in species mixtures, because a greater number of different forms of soil P were used compared to those present in monocultures (Zou, Binkley, and Caldwell 1995; Bu et al. 2020). Additionally, in forests, increased species richness can increase litter diversity, abundance (Huang et al. 2017) and decomposition (Lin et al. 2021), thus increasing the incorporation of organic matter in the soil and contributing to enriching it with nutrients (Gartner and Cardon 2004). Aboveground, species richness has been positively linked to the spatial complementarity of tree crowns (or “canopy packing”, Jucker, Bouriaud, & Coomes, 2015), that is, a more complete use of available space for the canopies resulting in an improved interception of incoming light. Such possible effects on a tree individual’s trait variation resulting from plant interactions should be greatest from directly neighbouring plants, and fade for more distant neighbours (Vogt, Murrell, and Stoll 2010; Mokany, Ash, and Roxburgh 2008a). Few studies have so far attempted to look specifically at the effects of the neighbouring species richness on intra-specific trait variation (but see Benavides, Valladares, Wirth, Müller, & Scherer-Lorenzen, 2019; Le Bagousse-Pinguet et al., 2015) and even less addressed its effect on intra-individual trait variation (but see Proß et al., 2021), preventing any general conclusion.

To investigate changes in functional traits in response to soil resource availability and species richness of the local neighbourhood, tree diversity experiments are an ideal framework, as species richness is manipulated while other macro-ecological conditions, as well as plant density and age, are constant. In addition, trees are long-lived organisms for which trait variation and adaptation to varying environmental conditions is particularly critical. In this study, carried out in a subtropical tree diversity experiment, we aimed at determining the shares of different levels of trait variation (between species, between individuals, within individuals) to the overall trait variation. In addition, focusing specifically on intra-individual trait variation, we investigated the influences of soil nutrient availability, the identity of a tree’s directly adjacent (i.e., ‘direct’) neighbour, as well as the diversity of its surrounding neighbourhood.

We hypothesized that (I) the largest proportion of total leaf trait variation is explained by differences between species, but intra-individual variation represents a considerable share of the remaining intra-specific part (which includes both variation between and within individuals). Further, we expected intra-individual leaf trait variation (II) to be positively related to soil nutrient availability and (III) to be greater for trees that have a different tree species as their direct neighbour (hetero-specific direct neighbouring tree), and to increase with increasing species diversity of the surrounding neighbourhood. Thereby, we assumed the effect of the direct neighbour to be more important than the surrounding neighbourhood. Finally, we hypothesized that (IV) the positive effects of soil nutrient availability, a hetero-specific direct neighbouring tree and the surrounding neighbourhood diversity amplify each other, that is, that the positive effect resulting from their interaction is greater than the sum of their respective positive effect.

## Materials and Methods

### 1. Study site

Our study was conducted in the BEF-China tree diversity experiment, which was set up in subtropical China near Xingangshan in the Jiangxi Province (29.08–29.11 N, 117.90-117.93 E). The mean annual temperature in Xingangshan is 16.7 °C and the mean annual precipitation 1821 mm (Yang et al. 2013). Our study took place in Site B, a part of the experiment established in 2010. Site B covers 20 ha consisting of 295 plots of 25.81 m x 25.81 m, from which we used 57 for this study. Each plot comprises 400 trees, planted in a regular grid, 1.29 m apart. The plots vary in their species richness, ranging from monocultures to mixtures of two, four, eight, or 16 native tree species (Bruelheide et al. 2014).

### 2. Experimental design

To differentiate between the effect of a tree’s direct neighbour species identity (either mono- or hetero-specific) and the species diversity of its surrounding neighbours, our study focuses on tree species pairs (i.e., pairs of two directly adjacent trees; TSPs) and the ten neighbours immediately surrounding the pairs (i.e., the TSP’s local neighbourhood). Although all trees are planted at equal distance from each other, we considered trees within a TSP ‘direct neighbours’, as leaf traits were measured for both trees of a TSP at the side of the crown where the two trees are in direct contact (i.e. interaction plane; Supp. Fig. S1).

We investigated 14 tree species belonging to four groups of four species each (two species are in two groups; see Supp. Table S1 & S2), corresponding to four different four-species mixtures. Within each group, we considered all possible TSP combinations, with four mono-specific (AA, BB, CC, DD) and six hetero-specific TSP combinations (AB, AC, AD, BC, BD, CD). We sampled these ten TSP combinations in four-, eight- and 16-species mixture plots. In two-species mixtures, we sampled all three possible TSP combinations (AA, BB, AB) as well as mono-specific TSPs in all monocultures. To have sufficient replication, the TSP combinations were repeated three times in two-species mixtures and monocultures. The study design encompassed 222 TSPs, of which 220 could be sampled, allowing the study of 440 trees in total.

### 3. Leaf traits

#### Leaf sampling

Our study considers nine leaf traits: specific leaf area (SLA), leaf dry matter content (LDMC), leaf carbon (C) and leaf nitrogen (N) contents, leaf carbon to nitrogen ratio (C:N), leaf magnesium (Mg), leaf calcium (Ca), leaf potassium (K) and leaf phosphorus (P) contents. To investigate them, leaves from 440 trees were sampled between mid-August and mid-September 2019. For each tree, we sampled leaves at the TSP interaction plane (Supp. Fig. S1), at two to five heights along the crown, depending on crown extent. Three fully developed leaves were sampled at each height. In total, this led to over 5,800 leaf samples from the TSPs.

Because of the high number of leaves collected and because leaf material of each TSP sample (i.e., the three leaves from the same height within a tree crown) would be insufficient to conduct all chemical analyses, we used optical spectroscopy to predict leaf traits of the TSP samples based on additional calibration samples (see below for details on spectral data acquisition and predictions). These calibration samples were collected at the same time as the TSP samples and for all species considered, but not directly from the TSPs themselves. Calibration samples were collected at multiple heights and positions within the crown of 130 randomly selected trees, representative of all species across all plot diversity levels. Each of the 252 calibration samples was composed of ca. 15 leaves, depending on leaf size, in order to have sufficient material for laboratory analyses. After leaf collection, TSP and calibration samples were transported in a water saturated and cooled atmosphere to the laboratory facilities, to be stored at 6-8 °C until being processed.

#### Leaf spectral data acquisition

We used reflectance spectroscopy, including visible and near-infrared wavelengths (NIRS), as a fast and cost-effective method (Serbin et al. 2014; Burnett et al. 2021; Trogisch et al. 2017) to predict leaf trait values from spectral data (Supp. Fig. S2). Spectral data of all leaves (i.e., TSP and calibration samples) were acquired on the sampling day, using an ASD FieldSpec4 Wide Resolution Field Spectroradiometer (Malvern Panalytical Ltd., Malvern, United Kingdom) with the RS³-Software, over a 350 nm to 2500 nm wavelength range. The device was regularly calibrated with a diffuse reflectance target (Spectralon, Labsphere, Durham, New Hampshire, USA) as white reference. Using a leaf contact probe, three repeated spectral measures were taken on the upper surface of each fresh leaf, with every spectral measurement consisting of ten internally averaged recordings of the device.

#### Laboratory determination of calibration samples’ leaf traits

In order to predict leaf traits of TSP samples from their spectral data, leaf traits were first measured directly from the calibration samples. Fresh leaves from the calibration samples were weighed before spectral data acquisition (see above). Leaf area was then measured with a flatbed scanner and the WinFolia software (Regent Instruments, Quebec, Canada). Afterwards, leaves were dried at 80 °C for 72 h before being weighed again, and SLA (leaf area / leaf dry mass) and LDMC (leaf dry mass / leaf fresh mass) were calculated (Pérez-Harguindeguy et al. 2013).

To quantify chemical leaf traits, dry leaves were ground in a ball mill (MM 400, Retsch, Germany) and the resulting leaf powder divided for the different analyses. Total C and N content, from which C:N was calculated, were determined gas-chromatographically with an elemental analyser (vario EL cube, Elementar, Hanau, Germany). Following an HNO3 digestion, the filtrate was used to measure leaf Mg, Ca and K content by atomic absorption spectrometry (ContrAA 300 AAS, Analytik Jena, Jena, Germany), and leaf P content with an acid molybdate spectrophotometric assay (see Supp. Table S3 for trait summary statistics).

#### Leaf trait prediction models

Using the calibration samples’ measured leaf traits matched with their respective spectral data, we fitted predictive models to apply to our TSP samples. Since several leaves were pooled for measuring the traits of each of the 252 calibration samples, we matched each calibration sample’s averaged leaf spectral data with its single measured trait value.

With this calibration dataset, we then fitted Partial Least Square Regression (PLSR) models for each trait using the non-linear iterative partial least square algorithm (NIPALS), validated with full cross-validation. The stability of the model was maximized and the most informative wavelength ranges identified according to Marten’s uncertainty test (Martens, 1999). The model’s number of latent factors was based on the amount of variance of the response variable explained by the model, so that a minimum of factors explained a maximum of variance, avoiding model overfitting. Different mathematical pre-treatments were applied to the averaged calibration samples’ spectra in order to reduce noise and highlight their most relevant features (Supp. Table S4). The PLSR model was then used to predict trait values from each spectral measure of our TSP leaf samples. Because we took three spectral measures per leaf, the three predictions per leaf were averaged after outlier identification (see below for details in Data analysis), resulting in one predicted trait value for each leaf (Supp. Table S3). The R-Square (R^2^) and root mean square error of the prediction (RMSE) were used to estimate robustness and performance of the models (Supp. Table S4). All spectral data analyses were performed with the Unscrambler X software (Version 10.5.1, CAMO Analytics, Oslo, Norway).

### 4. Soil data

#### Sampling

For each of the 220 TSPs, one bulk soil sample was taken at equal distance between both trees of a TSP, and one meter away from the interaction plane (Supp. Fig. S1). About 30 g of fresh soil were collected from the mineral layer, between 5 and 10 cm depth. Samples were kept cool at 4°C until being processed.

#### Laboratory analyses

To characterise soil nutrient availability, we measured soil P content, soil C:N and soil cation exchange capacity (CEC). The fresh soil samples were sieved to 2 mm. Soil P was measured photometrically using the Olsen method (Olsen 1954). To determine CEC, we performed a percolation with barium chloride. In the resulting percolate, the concentrations of Ca, Mg and K were analysed by atomic absorption spectrometry (ContrAA 300 AAS, Analytik Jena, Jena, Germany), and by measuring the pH of the percolate, we calculated the hydrogen concentration. Cation exchange capacity was then calculated as the sum of the ion equivalents of all cations measured (Ca, Mg, K and H). To determine total C and total N contents, the sieved soil was dried at 105 °C and milled to fine powder. The C and N contents were obtained by gas chromatography performed with an elemental analyser (vario EL cube, Elementar, Hanau, Germany), from which soil C:N was then calculated.

### 5. Data analysis

In order to determine the proportion of total trait variation explained by differences between species, between individuals within a species, and within individuals (hypothesis I), linear mixed-effects models were fitted for each trait (“lme” function in the R package “nlme”, Pinheiro et al., 2017). The response variable consisted of the predicted trait values for each repeated leaf spectral measure (i.e. three values for each leaf), and as fixed effect, only the intercept was included. As random term we included the leaf nested in sampling height (representing the intra-individual contribution to intra-specific trait variation; ITV_I_) nested in tree identity (representing the between-individual contribution to intra-specific trait variation; ITV_B_) nested in species identity (representing the between-species trait variation; BTV). The function “varcomp” from the R package “ape” (Paradis and Schliep 2019) was used to obtain the variance of each level of trait variation.

For hypotheses II, III and IV, we used Rao’s quadratic entropy (Rao’s Q; Botta-Dukát, 2005) to quantify intra-individual trait variation (ITV_I_). Rao’s Q was calculated with the “dbFD” function from the package “FD” (Laliberté, Legendre, and Shipley. 2014) for each trait of each tree, using trait values averaged at the leaf level. Considering the design of our study, weights of each sample (that is, of each leaf) within a tree individual were equal, and thus all set to one. This way, Rao’s Q gives the average Euclidian distance between the trait values of all leaves within one individual, as a measure of within-tree variation.

To exclude bias arising from differences in sample size per tree, we first fitted linear mixed-effects models (“lmer” function from R package “lmerTest”; Kuznetsova, Brockhoff, & Christensen, 2017) with Raós Q (i.e., ITV_I_) as response to the number of values used to calculate Rao’s Q. As random factors, TSP identity was nested in plot identity, and as a crossed random factor, species identity of the tree were added in the models. We extracted the residual variance of these models, and in a second step, used simple linear models to explain the residual variance with soil nutrient availability (CEC, soil C:N ratio, soil P content), TSP diversity (i.e., mono- or hetero-specific TSP partner) and neighbourhood diversity (i.e., Shannon Diversity Index of the trees directly surrounding the TSP) as fixed effects. The interactions of all fixed effects were added, except for interactions between soil variables. The Shannon Diversity Index, quantifying the species richness and evenness within a community, was calculated using the “vegan” package (Oksanen et al. 2020).

Before averaging the trait values predicted for each spectral measure taken (i.e., three per leaf), we excluded outlying predicted trait values (i.e., negative values, values exceeding a 5 % deviation from the range limits of calibration data, and values outside a 95 % confidence interval around the predicted values’ distribution). Before calculating Rao’s Q from the resulting averaged trait values at the leaf level, we also made sure that leaf trait values originated from at least two different sampling heights within a tree. If either tree of a TSP did not fulfil these criteria, i.e. had either only outlying trait values or all trait values came from one sampling height, we excluded the entire TSP. As a result, we included between 195 and 207 TSPs in the analyses, depending on the trait. Model assumptions were checked with the R package “performance” (Lüdecke et al. 2021). To correct for non-normality of the residuals, Raós Q values were log-transformed in all models. The full models were then simplified by stepwise removal of model terms based on Akaike’s information criterion (AIC, function “stepAIC” from package “MASS”; Venables & Ripley, 2002). All statistical analyses were carried out with R version 4.0.4 (R Core Team 2021).

## Results

### 1. Distribution of trait variation

Averaged across all traits, intra-specific trait variation (ITV_I_ and ITV_B_ taken together) represented the largest share of the total variation (43%; Fig 1). Specifically, for five out of the nine traits studied (SLA, leaf C:N, leaf N, leaf Mg and leaf Ca), the proportion of ITV was greater than that of BTV. Within ITV, intra-individual trait variation (ITV**_I_**, i.e. variation driven by differences between sampling heights) accounted on average for more than a quarter of the total variation (27%), and between-individual trait variation (ITV_B_) for around 16%. In particular, ITV**_I_** exceeded 25% for four traits (SLA, leaf C:N, leaf N, leaf K). For SLA and leaf C:N, ITV**_I_** was also greater than BTV (49 % vs. 38 % and 39 % vs. 28 % respectively).

**Figure 1:**
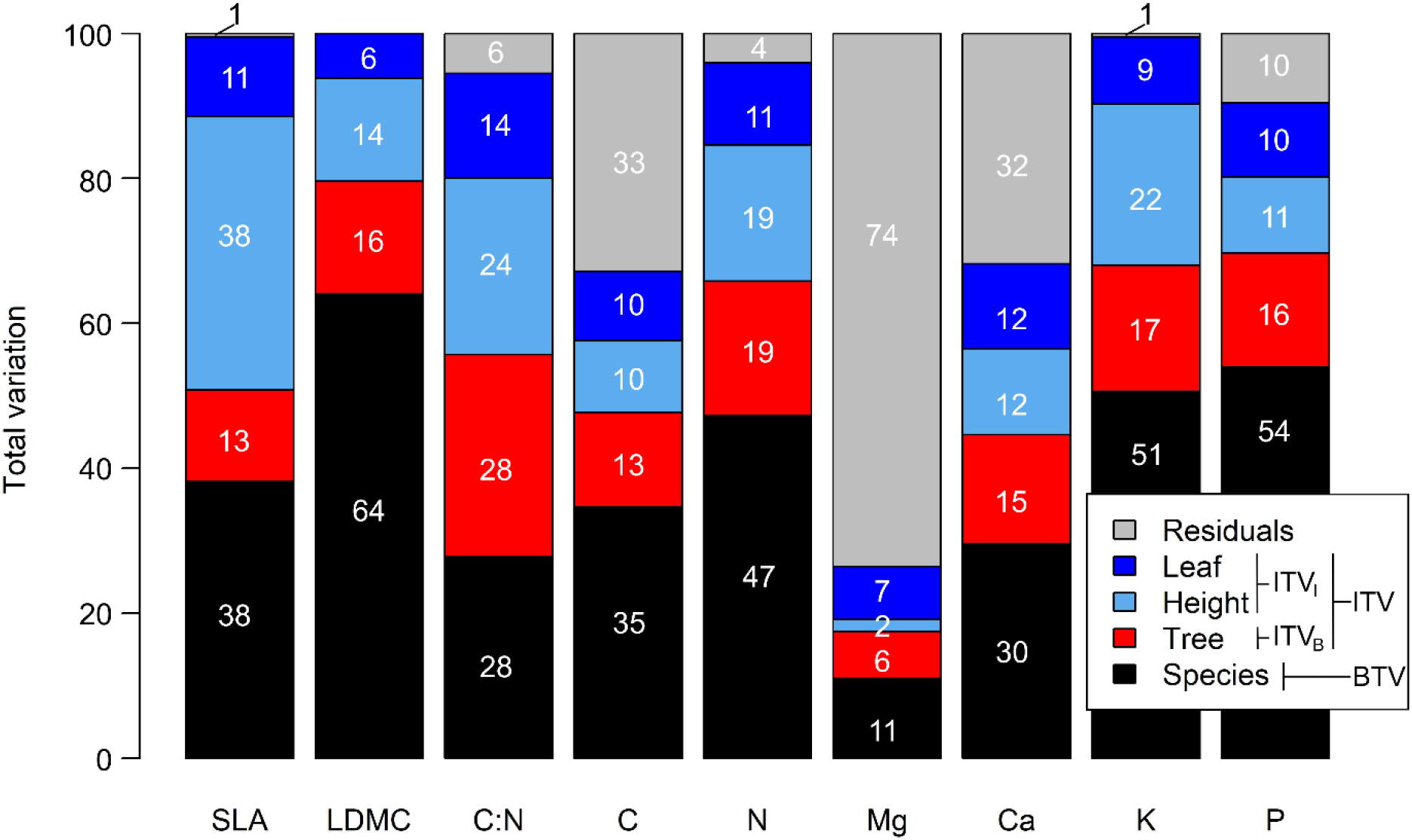
Partitioning of the variation in leaf traits at different scales: between species (BTV – between species trait variation) and within species (ITV – intra-specific trait variation). The latter separates into between trees trait variation (ITV_B_) and within tree trait variation (ITV_I_ – intra-individual variation including between heights and between leaves variation). Amount of variation is indicated in percent (%). Residual variation includes variation between repeated collection of leaf spectra (within-leaf variation) and error.

The proportions of BTV and ITV**_I_** differed greatly between traits, with values for BTV ranging from 11 % (leaf Mg) to 64 % (LDMC) of the total variation, and for ITV**_I_** between 9 % (leaf Mg) and 49 % (SLA). Also, ITV_B_ varied between traits, ranging from 6 % to 28 % (resp. leaf Mg and leaf C:N). For most traits, the proportion of the total variation which could not be explained by the scales studied (BTV, ITV_B_ or ITV**_I_**) was small (10% or less), except for leaf Ca (32%), leaf C (33%) and leaf Mg (74%).

### 2. Effects of chemical soil properties on intra-individual leaf trait variation

For three out of the nine leaf traits studied (SLA, leaf C:N, and leaf K), we found partly an increase of ITV**_I_** with higher nutrient availability in the soil, i.e. higher soil P content or lower soil C:N (Fig 2).

**Figure 2:**
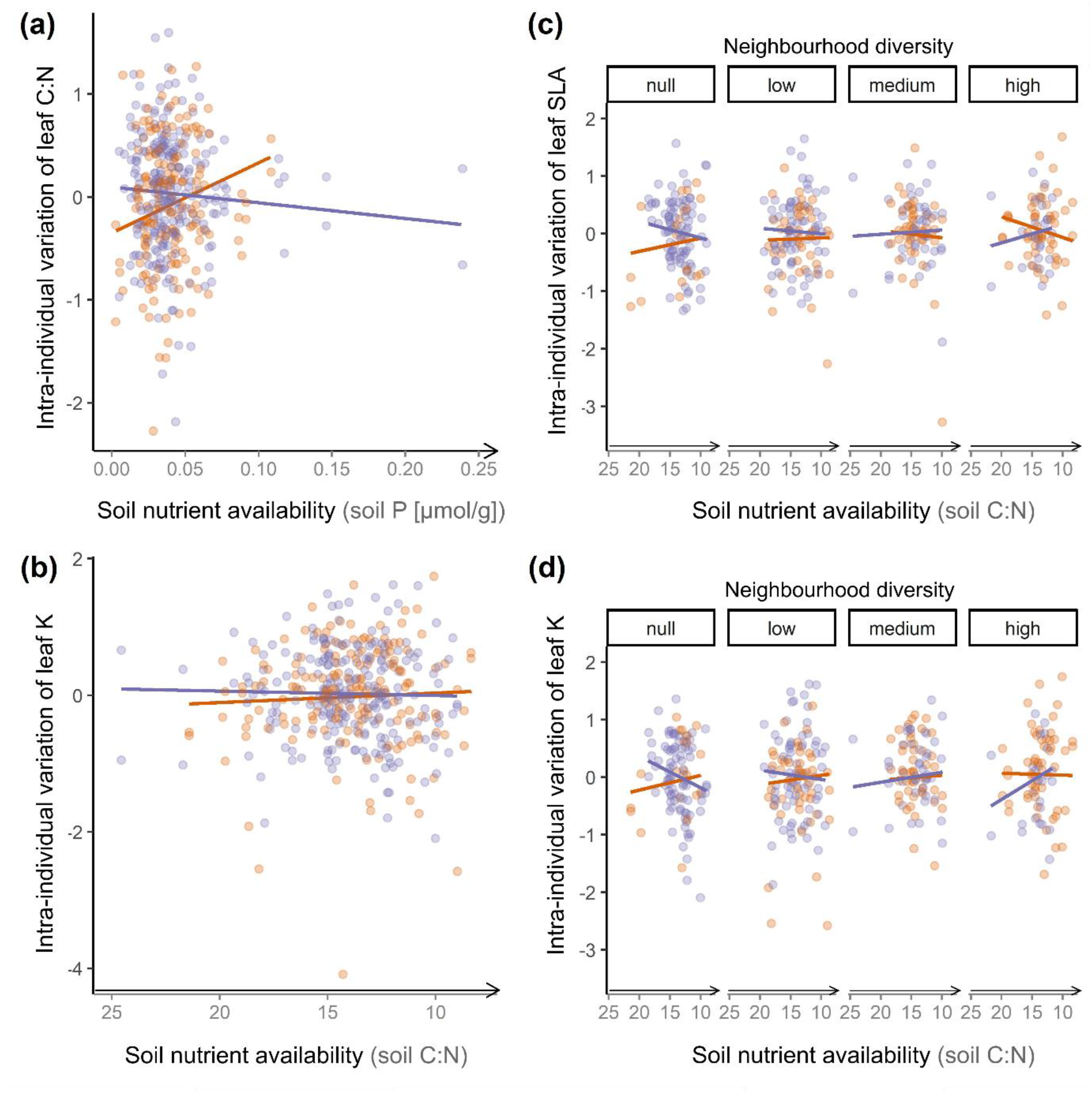
Interacting effects of soil nutrient content and species diversity on intra-individual leaf trait variation. Diversity of the tree species pair (orange line: hetero-specific direct neighbour; purple line: mono-specific direct neighbour) interact with the soil nutrient availability (soil P; soil C:N) and modify the intra-individual variation of leaf C:N **(a)** and leaf K **(b)**. Specific leaf area (SLA) **(c)** and leaf K **(d)** also vary depending on TSP diversity interacting with soil nutrient content, but also local neighbourhood diversity (Shannon Diversity Index of the ten trees surrounding a pair; null: SDI=0, low: 0<SDI≤0.8, medium: 0.8<SDI≤1.1, high: 1.1<SDI≤1.8). Orange represents hetero-specific TSPs and purple, mono-specific TSPs. Intra-individual trait variation is calculated as logarithm-transformed Rao’s Q value of each tree; therefore the scale of variation ranges from negative to positive values. Increasing soil nutrient availability corresponds to increasing soil P content or decreasing soil C:N, hence, soil C:N axis is shown in reverse order. Only significant effects are shown (Table 1).

**Table 1:** Model results analysing the interacting effects of neighbourhood diversity (Shannon Diversity Index of the ten trees surrounding a TSP), TSP diversity (mono- or hetero-specific direct neighbour) and soil nutrient variables (soil C:N, soil P and Cation Exchange Capacity) on the intra-individual variation of nine leaf traits (calculated as the log-transformed Rao’s Q value of each studied tree). Full models were specified to include all possible interactions, except for the interaction between several soil nutrient variables, and simplified by stepwise removal of terms based on model AIC. Significant effects (p<0.05) of the most parsimonious models are indicated in bold. Abbreviations: SLA: specific leaf area; LDMC: leaf dry matter content; C:N: leaf carbon to nitrogen ratio; C: leaf carbon content; N: leaf nitrogen content; Mg: leaf magnesium content; Ca: leaf calcium content; K: leaf potassium content; P: leaf phosphorus content; Neigh. div.: local neighbourhood diversity; TSP div.: diversity of the tree species pair; CEC: cation exchange capacity; soil C:N: soil carbon to nitrogen ratio; soil P: soil phosphorus content.

| Intra-individual leaf trait variation | Predictor | Estimate | Std. Error | F-value | p-value |
| --- | --- | --- | --- | --- | --- |
| SLA | Neigh. div. | -0.462 | 0.467 | 0.979 | 0.323 |
|  | TSP div. | -0.737 | 0.598 | 1.520 | 0.218 |
|  | Soil C/N | -0.032 | 0.033 | 0.982 | 0.322 |
|  | Neigh. div. x TSP div. | 1.124 | 0.663 | 2.876 | 0.091 |
|  | Neigh. div. x Soil C:N | 0.047 | 0.033 | 1.989 | 0.159 |
|  | TSP div. x Soil C:N | 0.071 | 0.043 | 2.733 | 0.099 |
|  | Neigh. div. x TSP div. x Soil C:N | -0.095 | 0.047 | 4.040 | **0.045** |
| LDMC | Neigh. div. | 0.247 | 0.131 | 3.536 | 0.061 |
|  | TSP div. | -0.005 | 0.243 | 0.000 | 0.985 |
|  | CEC | -0.048 | 0.028 | 2.953 | 0.086 |
|  | Neigh. div. x TSP div. | -0.330 | 0.163 | 4.122 | **0.043** |
|  | TSP div. x CEC | 0.057 | 0.032 | 3.168 | 0.076 |
| C:N | Neigh. div. | 0.526 | 0.266 | 3.926 | **0.048** |
|  | TSP div. | 0.853 | 0.300 | 8.061 | **0.005** |
|  | Soil P | 15.686 | 6.915 | 5.145 | **0.024** |
|  | Neigh. div. x TSP div. | -0.584 | 0.305 | 3.675 | 0.056 |
|  | Neigh. div. x Soil P | -12.740 | 6.610 | 3.715 | 0.055 |
|  | TSP div. x Soil P | -16.864 | 7.104 | 5.635 | **0.018** |
|  | Neigh. div. x TSP div. x Soil P | 12.237 | 7.362 | 2.763 | 0.097 |
| C | TSP div. | 0.083 | 0.043 | 3.715 | 0.055 |
| N | TSP div. | 0.209 | 0.136 | 2.339 | 0.127 |
|  | CEC | 0.016 | 0.020 | 0.640 | 0.424 |
|  | TSP div. x CEC | -0.029 | 0.023 | 1.644 | 0.200 |
| Mg | Neigh. div. | -0.470 | 0.320 | 2.156 | 0.143 |
|  | TSP div. | -0.315 | 0.408 | 0.594 | 0.441 |
|  | Soil C:N | -0.011 | 0.022 | 0.256 | 0.613 |
|  | Neigh. div. x TSP div. | 0.804 | 0.455 | 3.123 | 0.078 |
|  | Neigh. div. x Soil C:N | 0.026 | 0.023 | 1.296 | 0.256 |
|  | TSP div. x Soil C:N | 0.012 | 0.029 | 0.174 | 0.677 |
|  | Neigh. div. x TSP div. x Soil C:N | -0.047 | 0.032 | 2.108 | 0.147 |
| Ca | Neigh. div. | 0.054 | 0.080 | 0.446 | 0.505 |
|  | TSP div. | 0.046 | 0.094 | 0.239 | 0.625 |
|  | Neigh. div. x TSP div. | -0.087 | 0.099 | 0.771 | 0.381 |
| K | Neigh. div. | -0.227 | 0.549 | 0.171 | 0.680 |
|  | TSP div. | -1.322 | 0.704 | 3.532 | 0.061 |
|  | Soil C:N | -0.030 | 0.038 | 0.623 | 0.430 |
|  | Neigh. div. x TSP div. | 1.546 | 0.783 | 3.901 | **0.049** |
|  | Neigh. div. x Soil C:N | 0.023 | 0.039 | 0.344 | 0.558 |
|  | TSP div. x Soil C:N | 0.104 | 0.050 | 4.266 | **0.040** |
|  | Neigh. div. x TSP div. x Soil C:N | -0.119 | 0.056 | 4.542 | **0.034** |
| P | TSP div. | 0.670 | 0.326 | 4.237 | **0.040** |
|  | Soil C:N | 0.025 | 0.017 | 2.166 | 0.142 |
|  | TSP div. x Soil C:N | -0.042 | 0.023 | 3.305 | 0.070 |

Overall, leaf C:N variation increased with increasing soil P content, which was mainly driven by the steep increase of leaf C:N variation in hetero-specific TSPs, while leaf C:N variation in mono-specific TSPs decreased with increasing soil P content (Table 1; Fig. 2a). Similarly, leaf K of trees with a hetero-specific neighbour showed an increase in variation with increasing soil nutrient, while variation of trees within mono-specific TSPs decreased. However, mono- and hetero-specific TSPs were most similar in leaf K variation at highest nutrient availability, while leaf C:N variation was most similar for mono- and hetero-specific TSPs at the lowest nutrient availability (Table 1; Fig. 2a, with increasing soil P, and 2b, with decreasing soil C:N).

The described contrasting effect of soil C:N on leaf K variation in mono- and hetero-specific TSPs was not consistent across neighbourhood diversity levels. Rather, for leaf K as well as SLA in hetero-specific TSPs, the effect of higher soil nutrients (that is, lower soil C:N) on variation was positive at null and low neighbourhood diversity, slightly positive (leaf K) or negative (SLA) at medium neighbourhood diversity, and negative at high neighbourhood diversity. For mono-specific TSPs, we found the opposite pattern, with increasing soil nutrients having a negative effect on trait variation at null and low neighbourhood diversity and a positive effect at medium and high neighbourhood diversity (Fig. 2c and d).

### 3. Diversity effects on intra-individual leaf trait variation

The results showed evidence of both TSP diversity and neighbourhood diversity effects on ITV**_I_**. Trees in mono-specific TSPs displayed a larger variation of leaf P and leaf C:N than those growing in hetero-specific TSPs (Table 1; Fig. 3a and b).

**Figure 3:**
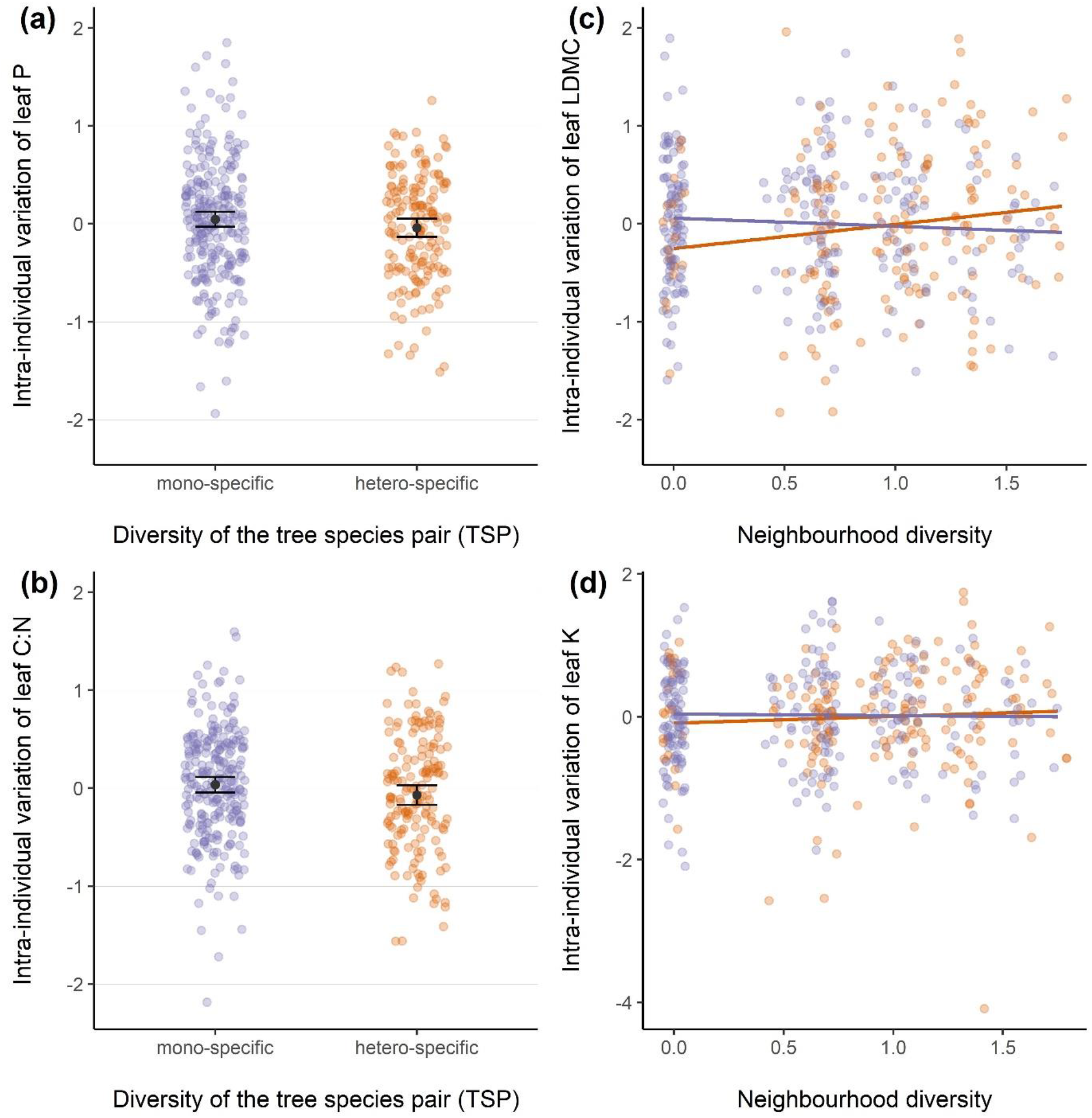
Effects of diversity on the intra-individual leaf trait variation. **(a)** and **(b)**: effect of the diversity of the tree species pair – mono- or hetero-specific direct neighbour – on leaf P and leaf C:N variation, respectively. Black dots indicate the mean of each group, and error bars two standard errors around the mean. The diversity of the tree species pair’s interaction with local neighbourhood diversity (calculated as the Shannon Diversity Index of the ten trees surrounding the TSP) modifies **(c)** leaf dry matter content (LDMC) variation, and **(d)** leaf C:N variation. Orange represents hetero-specific TSPs and purple, mono-specific TSPs. Intra-individual trait variation is the logarithm-transformed Rao’s Q value of each tree; therefore the scale of variation ranges from negative to positive values. Only significant effects are shown (Table 1).

While these effects seem to illustrate a negative relationship between diversity and trait variation, interaction between TSP and neighbourhood diversity showed that LDMC and leaf K variation decreased with neighbourhood diversity only for mono-specific TSPs, but increased for hetero-specific TSPs (Fig. 3c and d). For leaf C:N, we also found an overall slight increase of variation with neighbourhood diversity (Table 1). For four out of the nine leaf traits studied (leaf C, leaf N, leaf Mg and leaf Ca), no significant effects of soil nutrient availability and/or neighbourhood diversity on trait variation were found (Table 1).

## Discussion

Using an extensive dataset with over 5,800 leaf trait values collected in a large tree diversity experiment, our study is the first to consistently find, across a range of morphological and biochemical leaf traits, that the contribution of intra-specific trait variation to total trait variation amounts to a similar or even higher magnitude than the contribution of variation between species. Moreover, our results demonstrate that, within intra-specific trait variation, trait variation within individual trees holds a substantial share beside trait variation between individuals of the same species. Focusing on the intra-individual level, we show positive effects of soil nutrient conditions on trait variation. However, these were contingent on local species diversity.

### Intra-individual level: a substantial share of the total trait variation

Confirming our first hypothesis, we found that intra-individual variation (ITV_I_) represented on average over a quarter of the total variation, and was especially important in SLA and leaf C:N, where it exceeded the variation between species. Intra-specific variation, divided into variation between trees (ITV_B_) and within tree individuals (ITV_I_), was most often dominated by ITV_I_ or in similar proportions. This is in contrast to most of the few studies that included ITV_I_ in the comparison to higher levels of variation (e.g., Auger & Shipley, 2013; Hulshof & Swenson, 2010; Messier, McGill, & Lechowicz, 2010), as ITV_I_ was mostly found to explain the least amount of variation (but see Herrera, Medrano, & Bazaga, 2015). Between-species trait variation (BTV) displayed the largest share of variation, aligning with the findings of previous studies (Albert et al. 2010; de Bello et al. 2011; Siefert et al. 2015), but only when compared to the other levels of variation in isolation. In contrast, the total ITV added up to the largest proportion of variation for the majority of traits, suggesting that overlooking lower ecological levels (i.e. between and intra-individual) could lead to underestimating the share of intra-specific variation.

Even though largely consistent with the overall trend, the partition of variation showed different patterns for some traits. For example, SLA had an outstandingly large ITV_I_. Specific leaf area is known to respond strongly to local light conditions and hence to vary substantially in response to the light gradient of individual tree canopies (Niinemets, Keenan, and Hallik 2015). By adjusting the values of SLA, plants are able to optimise their light capture and thus their photosynthesis to adapt to light heterogeneity at the crown scale.

While most of the traits had a considerably larger share of ITV compared to BTV, the opposite occurred for LDMC, leaf K and leaf P. In particular, we found the largest proportion of BTV for LDMC. Considering the limited spatial extent of the study site, we can assume that the environmental variables, including the location’s water conditions, are relatively homogeneous across the site. As LDMC is primarily dependent on water availability (Niinemets 2001), this could explain a lesser importance of ITV_B_ and ITV_I_. In addition, contrary to other abiotic conditions (e.g., light), water availability does not present such strong variability within the tree crown, therefore contributing to a lower ITV_I_.

Hence, differences between traits regarding how their variation is partitioned could not be assigned, for example, to the trait type (morphological or chemical) or the leaf economics spectrum (i.e. acquisitive or conservative growth strategy; Reich 2014), but seem to arise from the spatial variability of the respective trait’s most important environmental driver (e.g., light, water availability etc.).

### Effects of soil nutrient availability on intra-individual trait variation

For three out of the nine leaf traits studied (SLA, leaf C:N and leaf K), we found an increase of ITV_I_ with higher nutrient availability in the soil, however contingent to species diversity conditions. Previous works showed that trait variation can be favoured by better soil conditions at the intra-specific level (Lemke et al. 2015; Helsen et al. 2017). For example, Lemke *et al*. (2012) reported for five herbaceous species in Germany an overall increase of intra-specific trait variation of both vegetative and reproductive traits with increasing soil P and soil N concentration. Our results suggest that better soil nutrient availability can promote trait variation not only at the intra-specific level, but also at the intra-individual level. However, apart from the positive effect of soil P concentration on leaf C:N variation, none of the soil effects was consistent across neighbourhood diversity levels, and no soil effect was independent of TSP diversity.

Previous studies reported cases of higher soil nutrient availability and plant nutrient concentrations in more diverse neighbourhoods in comparison to less diverse ones (Zak et al. 2003; Fargione et al. 2007; Dybzinski et al. 2008). This pattern has been suggested to result primarily from a) resource-use complementarity (Ashton et al. 2010; McKane et al. 2002), b) increased litter diversity, abundance and decomposability (Huang et al. 2017; Lin et al. 2021), and c) increased nutrient uptake arising from of higher microbial diversity. A greater diversity of fungi and bacteria has been shown to improve ecosystem soil functions, for example by increasing plant nutrient uptake and litter decomposition (Wagg et al. 2019). This was also observed for our study site in BEF-China, where Singavarapu *et al*. (2021) found that in species mixtures, fungal and bacterial communities exhibited higher species richness. However, in the results of the present study, the potentially positive effects of diversity on soil nutrients were not consistently reflected by intra-individual trait variation. Instead, we also found a decrease of intra-individual trait variation with increasing soil nutrient availability under multiple species diversity conditions. Depending on the leaf trait, this decrease was observed either generally for trees in mono-specific TSPs (leaf C:N), or in particular for trees in mono-specific TSPs at null or low neighbourhood diversity, and for trees in hetero-specific TSPs at medium and high neighbourhood diversity (SLA, leaf K).

Indeed, for both traits responding to the interaction of soil nutrient availability and diversity (SLA, leaf K), the slope of the soil-trait variation relationship described a hump-shape in response to increasing diversity (i.e., taking into account both TSP and neighbourhood diversity together; Fig. 4), peaking at medium diversity. Hence, at both ends of the diversity gradient, a low diversity environment (null or low neighbourhood diversity for mono-specific TSPs) and a highly diverse environment (medium or high neighbourhood diversity for hetero-specific TSPs) had a similarly negative effect on the soil-trait variation relationship. Inversely, the greatest increase of variation with soil nutrients occurred in moderately diverse environments, that is, at low neighbourhood diversity for hetero-specific TSPs and at medium neighbourhood diversity for mono-specific TSPs.

**Figure 4:**
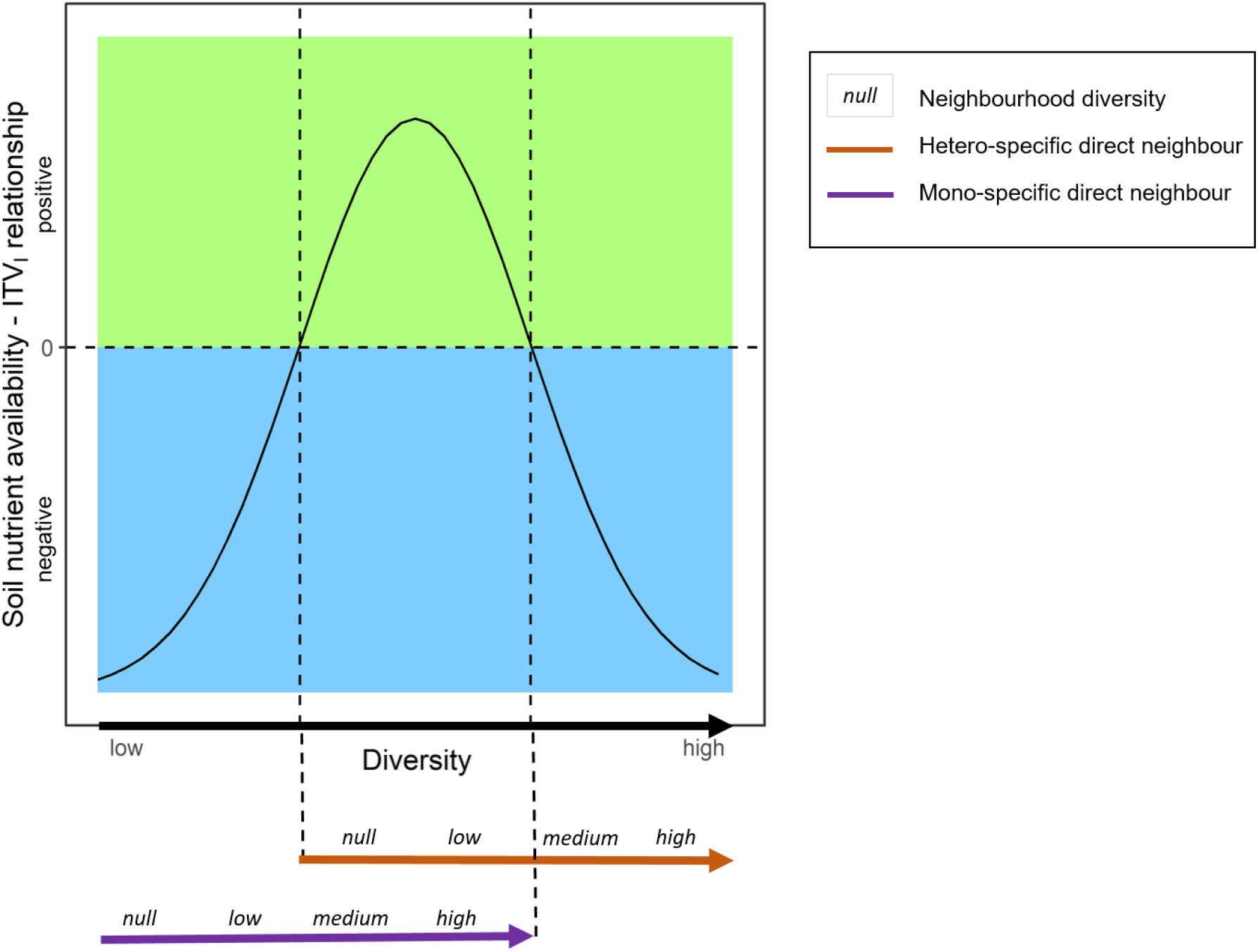
Conceptual representation of the effect of diversity on the relationship between soil nutrient availability and intra-individual trait variation (slope of the regression; represented by the black curve), derived from the response of SLA and leaf K to the drivers (see Fig. 2 c and d). Here, “diversity” considers jointly the diversity of the tree species pair (TSP), that is, having a mono or hetero-specific direct neighbour, and the local neighbourhood diversity, calculated as the Shannon Diversity Index of the ten trees surrounding a TSP. Green area represents an increase of variation with higher soil nutrient availability and blue area, a decrease of variation. The orange and purple arrows indicate the breaking down of diversity into TSP (hetero- or mono-specific direct neighbour) and neighbourhood (italic font levels) components.

The fact that we found negative relationships between soil nutrient availability and intra-individual trait variation at both ends of the diversity gradient for these two traits suggests that different mechanisms are involved. A possible explanation is that in conditions with very low diversity, intra-specific competition from either the mono-specific direct neighbour or surrounding mono-specific neighbours is the main mechanism at play, making trait variation a means to avoid competition. With soil nutrient availability increasing in such low-diversity environments, the opportunity for better nutrient uptake might alleviate the necessity to vary (Forrester 2014). On the other end of the gradient, high diversity might result in comparably negative effects for soil-trait variation relationship. Indeed, species richness has previously been shown to reach a threshold beyond which positive diversity effects seem to be overtaken by inter-specific competition, as the chance to encounter a strong competitor increases with the number of neighbouring species (Davrinche and Haider 2021). As under low-diversity conditions, greater soil nutrient availability in highly diverse environments might improve the growing conditions of hetero-specific TSPs to such an extent that the need for variation as a way to mitigate competition is remarkably reduced. Finally, moderately diverse conditions might result in the dominance of the positive effects of diversity (i.e., resource-use complementarity; litter diversity, abundance and decomposability; microbial diversity), thus reducing competition. Consequently, these positive diversity effects together with increased soil nutrients availability would enable a greater exploitation of resources, and thus the capacity for more trait variation. Hence, for specific traits, moderate species diversity could increase the possibility to maximise a tree’s local adaptation to changing environmental conditions.

### Effects of diversity on intra-individual trait variation

We also found effects of diversity independent of soil conditions: for leaf P and leaf C:N, mono-specific TSPs varied on average more than hetero-specific TSPs, contrary to our expectation (hypothesis III). While the overall response of leaf C:N variation was an increase with neighbourhood diversity, the TSP and neighbourhood diversity interaction resulted in an increase in variation with neighbourhood richness for hetero-specific TSPs but a decrease for mono-specifics for LDMC and leaf K. This increase in hetero-specific TSPs’ trait variation at higher neighbourhood diversity can be explained by the effect of diversity improving the use of belowground resources (i.e., resource-use complementarity, as described above), increasing resource uptake and hence enabling for more variation in trait values. In addition to the belowground partition of resources, aboveground spatial niche complementarity might also occur with increasing neighbourhood diversity. In a study also conducted within trees’ local neighbourhoods in the BEF-China experiment, Kunz et al. (2019) found crown packing to be denser in mixtures compared to monocultures, that is, that canopy space was used more efficiently. This might increase the gradient of light within a tree crown and, consequently, the variation of leaf traits.

In contrast, the decrease of intra-individual variation for LDMC and leaf K with increasing neighbourhood diversity for trees within mono-specific TSPs could be related to the greatest influence of the direct neighbour in comparison to more distant neighbours, in line with our assumption (hypothesis III). Indeed, having a direct neighbour from the same species implies a strong effect of intra-specific competition for trees in mono-specific TSPs. Hence, while increasing neighbourhood diversity might enable trees in hetero-specific TSPs to vary more by increasing the resource uptake and consequently the possibility to vary, it might also reduce their need for variation by alleviating the intra-specific competition for trees within mono-specific TSPs.

### Effects of interacting soil nutrient availability and diversity on intra-individual trait variation

Considering the observed interdependency of the effects of diversity and soil nutrient availability, we could not unequivocally refute or confirm our hypotheses II and III, assuming general positive effects of both drivers of intra-individual trait variation.

However, we found partial support for our fourth hypothesis: although there was no amplification of soil and diversity effects, intermediate species diversity ensured positive soil effects. Unexpectedly, both high and low extremes of species diversity reversed these effects, potentially revealing different processes behind what drives intra-individual trait variation. This emphasizes that the drivers of trait variation should not be considered in isolation, but are environment-dependent. Such an interplay is well-known, for example, from diversity-productivity relationships, which depends on environmental harshness (Mulder, Uliassi, & Doak, 2001), but has not been demonstrated for trait variation, and in particular not at the intra-individual scale.

Overall, our results strongly support the pledge for the inclusion of trait information below the species scale, when associating traits with ecosystem functions (Albert et al. 2010; Messier, McGill, and Lechowicz 2010; de Bello et al. 2011; Siefert et al. 2015; Hulshof and Swenson 2010; Kafuti et al. 2020). In particular, we could demonstrate for multiple traits the great share of intra-individual trait variation, which has not been found in such a consistent way and in a comparable extent so far. Regarding the abiotic and biotic drivers of intra-individual trait variation, our study is just a starting point. Even though the systematic exploration of multiple environmental drivers’ interacting effects on individual’s traits variation may be a challenge, knowledge about what drives and limits intra-individual trait variation will be crucial to understand local adaptations of plants to their complex environments. This will enable us to understand the potential of trees’ responses to changing environments due to, for example, global change, ultimately providing guidance for the conservation and rebuilding of sustainable forests.

## Supporting information

Supporting Information - Davrinche et al.

## Supporting Information

**Fig. S1:** Sampling design of leaves and soil material on Site B of BEF-China.

**Fig. S2:** Workflow for the application of spectral methods to leaf traits prediction.

**Table S1:** List of species included in the study.

**Table S2:** Broken-stick design of Site B of BEF-China.

**Table S3:** Summary statistics of leaf traits datasets.

**Table S4:** Parameters of the predictive models based on calibration samples’ spectral data.

## Acknowledgments

The study was supported by the International Research Training Group TreeDì jointly funded by the Deutsche Forschungsgemeinschaft (DFG, German Research Foundation) – 319936945/GRK2324 and the University of the Chinese Academy of Sciences (UCAS). We thank the local Xingangshan workers for maintenance of the field site and assisting in data collection, and the MLU student helpers for their help with processing the samples, as well as Yang Bo, Shan Li and Yuxi Xue for providing logistic assistance on site. SH, WSH, HB and AD conceived the ideas and designed methodology; AD and AB collected the data; AD and AB analysed the data, with support from SH, WSH, GA and HB; AD, AB and SH led the writing of the manuscript. All authors contributed critically to the drafts.

