## Supporting Information - Davrinche et al. for "High within-tree leaf trait variation and its response to species diversity and soil nutrients"

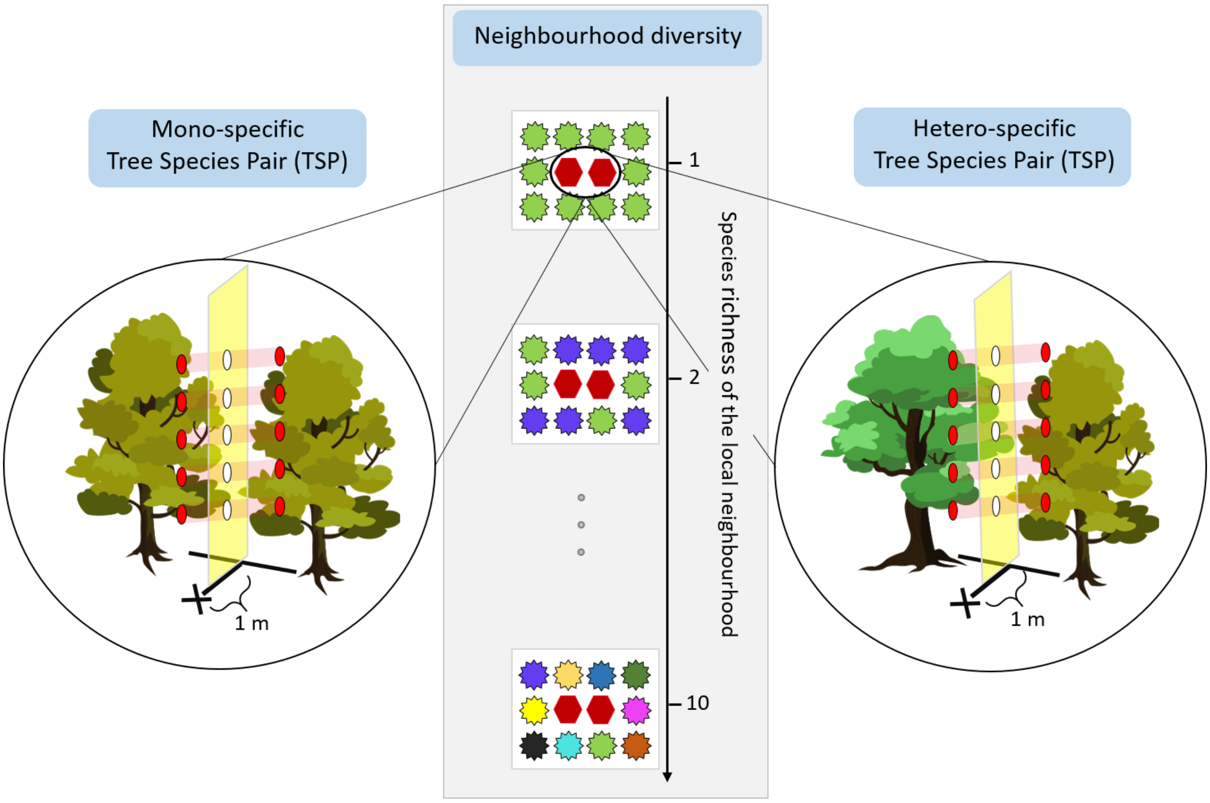


**Fig. S1:** Leaf and soil sampling design. Across Site B of the BEF-China experiment, we visited 220 Tree Species Pairs (TSP) in 57 plots. Around the TSP, the ten neighbouring trees belong to a gradient from one to ten different species (however, when accounting for tree mortality, this gradient is reduced to a maximum of six different species surrounding the TSP). Toothed shapes represent trees, red hexagons represent TSP partners and different colours represent different species. For each TSP, a soil sample was collected one meter away from the interaction plane of the two trees (black cross). For each tree, two to five samples (red dots) composed of three leaves each were taken along the crown, on the interaction plane between the two TSP partners. Adapted from Davrinche & Haider (2021).

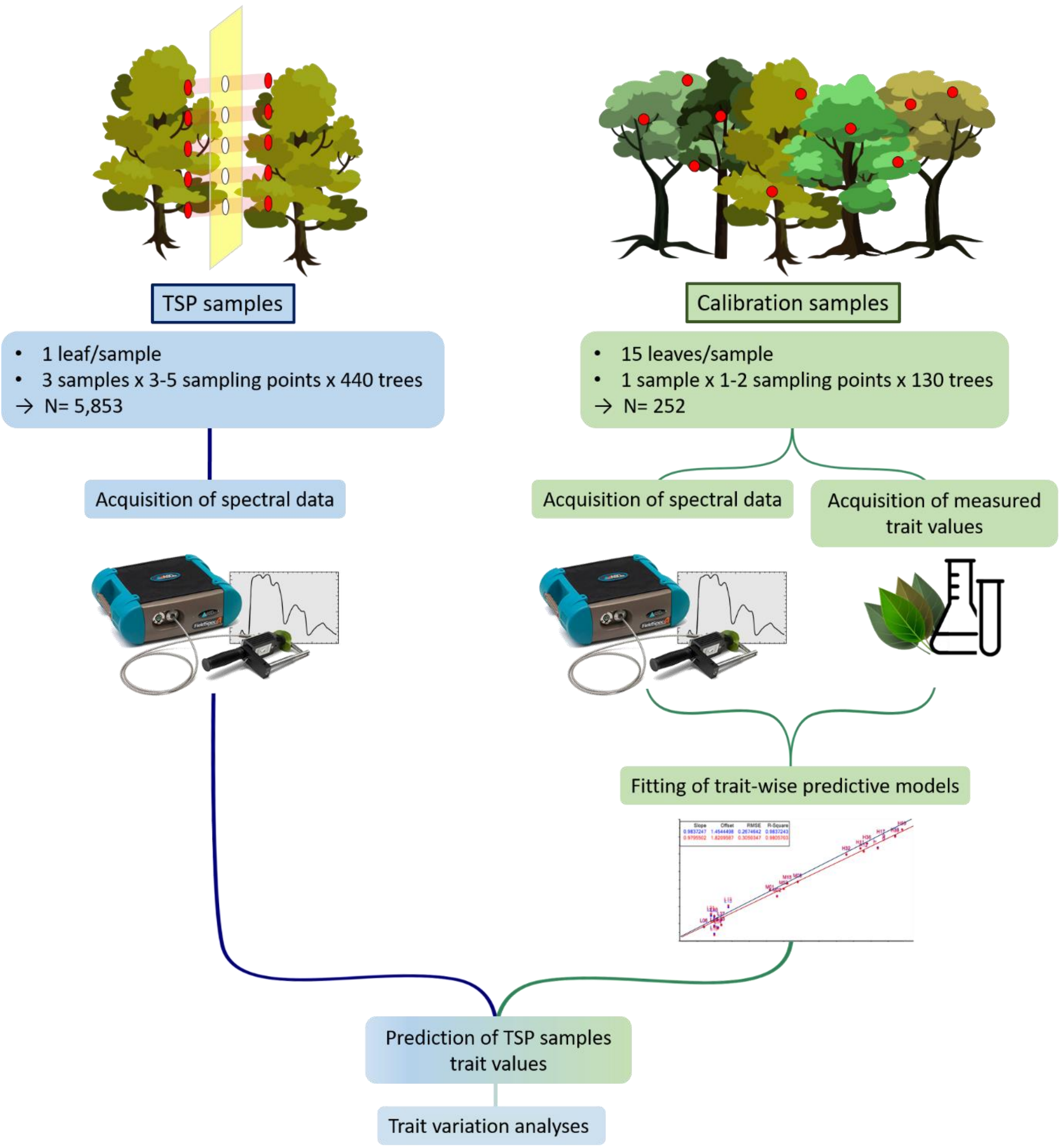


**Fig. S2:** Workflow for the application of spectral methods to leaf traits prediction. **Right side:** Samples from the tree species pairs (TSP) are collected, according to Fig. S1. Spectral data is acquired from the leaf samples of the TSP. **Left side:** In parallel, calibration samples are collected on randomly selected trees representative of the species and diversity levels used in the study. Both spectral data and measured values are acquired from calibration samples. The resulting dataset is used for creation and validation of predictive models for each trait, using partial least square regression. **Middle:** Predictive models are applied to the spectral data of TSP samples, to predict TSP samples' trait values. Those trait values are then used in the subsequent analyses, either at the spectral measurement level (three per leaf) for variance partitioning, or averaged at the leaf level for the calculation of intra-individual variation (as Rao's Q of each individual tree). Adapted from Burnett *et al*. (2021).

**Table S1:** List of tree species included in the study. Nomenclature according to The Flora of China (http://flora.huh.harvard.edu/china).

| Species name | Abbreviation | Author | Family |
|---|---|---|---|
| *Ailanthus altissima* | Aalt | (Miller) Swingle | Simaroubaceae |
| *Alniphyllum fortunei* | Afor | (Hemsley) Makino | Styracaceae |
| *Betula luminifera* | Blum | Winkler | Betulaceae |
| *Castanopsis fargesii* | Cfar | Franchet | Fagaceae |
| *Castanopsis sclerophylla* | Cscl | (Lindley & Paxton) Schottky | Fagaceae |
| *Cinnamomum camphora* | Ccam | (Linnaeus) Presl | Lauraceae |
| *Cyclobalanopsis glauca* | Cgla | (Thunberg) Oersted | Fagaceae |
| *Daphniphyllum oldhamii* | Dold | (Hemsley) Rosenthal | Daphniphyllaceae |
| *Elaeocarpus chinensis* | Echi | (Gardner & Champion) Hooker ex Bentham | Elaeocarpaceae |
| *Machillus leptophylla* | Mlep | Handel-Mazzetti | Lauraceae |
| *Machillus thunbergii* | Mthu | Siebold & Zuccarini | Lauraceae |
| *Manglietia fordiana* | Mfor | Oliver | Magnoliaceae |
| *Quercus phillyreoides* | Qphi | Gray | Fagaceae |
| *Schima superba* | Ssup | Gardner & Champion | Theaceae |

**Table S2:** Tree species pairs (TSPs) combinations within the broken-stick design of the BEF-China experiment (*sensu* Bruelheide et al., 2014). Abbreviations refer to species names in Table S1. Each cell of the 'Species combinations' column represents a group of species (including 1, 2, 4, 8 or 14 species). While the design originally counted 16 species, two of them did not have sufficient individuals to establish the required number of TSP, and were hence replaced by the repetition of two other species (*Alniphillum fortunei* (Afor) and *Castanopsis fargesii* (Cfar)). Each species occurs at each plot species richness level, and is studied in a combination with itself (mono-specific TSP) or another species from the same four-species group (hetero-specific TSP). For example, *Machilus leptophylla* (Mlep) and *Machilus thunbergii* (Mthu) can form a pair in a four-species mixture, but not in a two-species mixture. In order to keep the number of TSPs manageable, for eight- and 14-species mixtures, we used the same groups of species as in the four-species mixtures. According to all the possible combinations within groups of one, two or four species and their respective replication, the total number of tree species pairs (TSPs) sums up to 222 from which 220 were possible to sample for this study. Adapted from Bruelheide *et al.* (2014).

<table>
<tr><th>Plot species richness</th><th colspan="16">Species combinations within the BEF-China broken-stick design (Site B)</th><th>Theoretical number of combinations</th><th>Theoretical number of replication</th><th>Realised number of combinations</th><th>Total number of realised TSPs</th></tr>
<tr><td>14</td><td colspan="16">Afor Mlep Mthu Echi Blum Cfar Mfor Qphi Cgla Ssup Ccam Dold Cfar Cscl Aalt Afor</td><td>38</td><td>1</td><td>37</td><td>40</td></tr>
<tr><td>8</td><td colspan="8">Afor Mlep Mthu Echi Blum Cfar Mfor Qphi</td><td colspan="8">Cgla Ssup Ccam Dold Cfar Cscl Aalt Afor</td><td>38</td><td>1</td><td>32</td><td>33</td></tr>
<tr><td>4</td><td colspan="4">Afor Mlep Mthu Echi</td><td colspan="4">Blum Cfar Mfor Qphi</td><td colspan="4">Cgla Ssup Ccam Dold</td><td colspan="4">Cfar Cscl Aalt Afor</td><td>38</td><td>1</td><td>36</td><td>38</td></tr>
<tr><td>2</td><td colspan="2">Afor Mlep</td><td colspan="2">Mthu Echi</td><td colspan="2">Blum Cfar</td><td colspan="2">Mfor Qphi</td><td colspan="2">Cgla Ssup</td><td colspan="2">Ccam Dold</td><td colspan="2">Cfar Cscl</td><td colspan="2">Aalt Afor</td><td>22</td><td>3</td><td>22</td><td>68</td></tr>
<tr><td>1</td><td>Afor</td><td>Mlep</td><td>Mthu</td><td>Echi</td><td>Blum</td><td>Cfar</td><td>Mfor</td><td>Qphi</td><td>Cgla</td><td>Ssup</td><td>Ccam</td><td>Dold</td><td>Cfar</td><td>Cscl</td><td>Aalt</td><td>Afor</td><td>14</td><td>3</td><td>14</td><td>41</td></tr>
<tr><td></td><td colspan="16"></td><td></td><td></td><td></td><td>220</td></tr>
</table>

**Table S3:** Summary statistics of the leaf traits datasets. Calibration samples are composed of ca. 15 leaves each, collected from 130 randomly selected trees at the experimental site, across all species and diversity levels. Leaf trait values of calibration samples result from direct measurements in the laboratory. Leaf trait values of tree species pairs (TSP) samples (one leaf per sample) were predicted from their spectral data and the trait-wise prediction models built from the calibration dataset (Fig. S2). The TSP samples' values presented are the ones used to calculate intra-individual variation (Rao's Q), i.e., the predicted trait values after outlier identification and averaging the three trait predictions for each spectral measure at the leaf level (see Methods – Data analysis for details).

| | Trait | Unit | Nb. of samples | min | max | median | mean |
|---|---|---|---|---|---|---|---|
| Calibration samples (ca. 15 leaves per sample) | SLA | $mm^2/mg$ | 250 | 52.19 | 316.62 | 89.55 | 97.32 |
| | LDMC | mg/g | 252 | 253.70 | 650.70 | 442.90 | 443.70 |
| | CN | g/g | 248 | 13.65 | 53.82 | 33.12 | 33.33 |
| | C | % | 248 | 42.79 | 51.12 | 47.78 | 47.73 |
| | N | % | 249 | 0.92 | 3.48 | 1.43 | 1.50 |
| | Mg | mg/g | 250 | 0.40 | 7.03 | 1.43 | 1.73 |
| | Ca | mg/g | 250 | 0.83 | 19.40 | 2.99 | 3.95 |
| | K | mg/g | 250 | 1.85 | 17.67 | 6.89 | 7.47 |
| | P | mg/g | 250 | 0.06 | 1.72 | 0.59 | 0.66 |
| | | | Total = 252 | | | | |
| TSP samples (one leaf per sample) | SLA | $mm^2/mg$ | 5487 | 38.98 | 161.73 | 93.98 | 95.59 |
| | LDMC | mg/g | 5615 | 297.30 | 569.60 | 433.50 | 433.40 |
| | CN | g/g | 5638 | 20.33 | 47.49 | 33.93 | 33.83 |
| | C | % | 5677 | 44.44 | 50.49 | 47.50 | 47.48 |
| | N | % | 5634 | 0.92 | 2.11 | 1.50 | 1.51 |
| | Mg | mg/g | 5793 | 0.07 | 4.74 | 2.09 | 2.14 |
| | Ca | mg/g | 5741 | 0.03 | 10.57 | 4.84 | 4.99 |
| | K | mg/g | 5592 | 3.04 | 12.14 | 7.52 | 7.60 |
| | P | mg/g | 5611 | 0.23 | 1.08 | 0.62 | 0.64 |
| | | | Total = 5853 | | | | |

**Table S4**: Functional leaf traits analysed and their Partial Least Square Regression models (PLSR) characteristics, with accuracy of the models evaluated by their Root Mean Square Error (RMSE) and coefficient of determination ($R^2$). Mathematical transformations applied to the spectra before fitting the PLS models are indicated as pre-treatments. PLSR models are fitted on the calibration data described in Table S3.

| Leaf trait | Abbr. | Growth strategy | Unit | $R^2$ | Number of factors in the PLSR model | RMSE | Pre-treatment of spectra |
|---|---|---|---|---|---|---|---|
| Specific leaf area | SLA | acquisitive | $mm^2/mg$ | 89.88 | 7 | 9.35 | Standard normal variate; Savitzky-Golay 2nd derivative |
| Leaf dry matter content | LDMC | conservative | mg/g | 88.99 | 5 | 22.67 | Normalisation |
| Carbon to nitrogen ratio | C:N | conservative | g/g | 78.62 | 9 | 2.86 | Normalisation; Savitzky-Golay 2nd derivative |
| Carbon content | C | conservative | % | 78.62 | 8 | 0.69 | Normalisation; Savitzky-Golay 2nd derivative |
| Nitrogen content | N | acquisitive | % | 75.97 | 8 | 0.13 | Standard normal variate; Savitzky-Golay 2nd derivative |
| Magnesium content | Mg | acquisitive | mg/g | 60.46 | 8 | 0.65 | Standard normal variate; Savitzky-Golay 2nd derivative |
| Calcium content | Ca | acquisitive | mg/g | 58.84 | 8 | 1.71 | Savitzky-Golay 2nd derivative |
| Potassium content | K | acquisitive | mg/g | 75.97 | 2 | 2.08 | Normalisation |
| Phosphorus content | P | acquisitive | mg/g | 64.06 | 8 | 0.16 | Standard normal variate; Savitzky-Golay 2nd derivative |
